# Stopping vs Resting state during motor imagery paradigm

**DOI:** 10.1101/2021.06.15.448360

**Authors:** Bastien Orset, Kyuhwa Lee, Ricardo Chavarriaga, José del. R Millán

## Abstract

Current non-invasive Brain Machine interfaces commonly rely on the decoding of sustained motor imagery activity (MI). This approach enables a user to control brain-actuated devices by triggering predetermined motor actions. One major drawback of such strategy is that users are not trained to stop their actions. Indeed, the termination process involved in BMI is poorly understood with most of the studies assuming that the end of an MI action is similar to the resting state. Here we hypothesize that the process of stopping MI (MI termination) and resting state are two different processes that should be decoded independently due to the exhibition of different neural pattens. We compared the detection of both states transitions of an imagined movement, i.e. rest-to-movement (onset) and movement-to-rest (offset). Our results shows that both decoders show significant differences in term of performances and latency (N=17 Subjects) with the offset decoder able to detect faster and better MI termination. While studying this difference, we found that the offset decoder is primarily based on the use of features in Beta band which appears earlier. Based on this finding, we also proposed a Random Forrest based decoder which enable to distinguish three classes (MI, MI termination and REST).

## I. Introduction

A Brain-Machine Interface (BMI) aims at providing communication and control pathways for people with motor disabilities [1]. This is made possible by enabling users to control external devices and interact with their environment while bypassing the normal neuromuscular pathways. Primarily based on electroencephalogram (EEG), non-invasive BMIs are presented as great instruments to be used in novel forms of therapy such as for stroke rehabilitation [2]–[4].

EEG-BMIs are usually based on a well-known paradigm called Motor imagery (MI) which is the process where a person is asked to mentally rehearse a given motor activity without any over motor output [5]. While performing MI tasks, changes can be observed in the brain with respect to a resting state. These changes are usually referred to as ERDs (Event-Related Desynchronization) and are characterized by a decrease of power generally observed in μ [8-12] Hz and β [13-30] Hz bands [6], [7]. For such types of BMIs, users generally follow a training procedure where they are instructed either to perform MI or to not perform any task (resting state, REST). This paradigm is known as a Go/No-Go (GNG) task [3]. Similarly, this paradigm can also be performed sequentially with a person firstly instructed to be resting followed by an MI task (rest-to-movement transition, onset) [8].

By exploiting such patterns, a BMI can trigger predefined actions and notably trigger the activations of BMI effectors such as exoskeleton or functional electrical stimulation (FES). However, a major drawback of such decoding strategy is that users are not trained to stop their actions. Indeed, the MI termination process involved in BMI is poorly understood, most of the studies assumed that the end of an MI action is similar to the resting state, therefore, assuming that it corresponds to the disappearance of MI patterns [9].

Although this assumption is widely used in the BMI field, the existence of a particular neurophysiological signature observed after movement termination is however well known. This signature can be characterized by an increase of power (event-related synchronization, ERS) induced in the β band. Such synchronization, often called β rebound, can last for about a second. Although the role of β rebound is still under debate, it is currently thought to have a function of inhibition of the motor cortex by somatosensory processing [10], [11]. Oscillatory activity in the β band has been also linked to an active process to promote the existing motor set aiming to maintain the current sensorimotor or cognitive state (i.e., status quo) [12]. Such synchronizations can be explained by an increase of rhythmic activity paradoxically due to a decrease of the excitability of cortical neurons or inhibited cortical neurons [13], [14]. Importantly the presence of β rebound doesn’t require any muscle activation since it was also reported after an imaginary movement (MI task) [15]. When performing hand-related motor tasks, ERS can mainly be observed in the contralateral hand representation area. In the β band, this synchronization can also be seen in the supplementary motor area (SMA) located in mid-central areas of the brain with slightly higher frequencies and an earlier onset compared to the contralateral ERS [16], [17].

Neuroimaging studies have also investigated the difference in the motor response inhibition when performing GNG task or Stop-Signal Task (SST) with the last one instructing participants to respond as fast as possible to a stimulus (go trial) but to cancel any response when a stop signal is presented (stop signal). Comparing both tasks, they found notably that these tasks were engaging overlapping but distinct neural circuits with GNG engaging more the fronto-parietal control network while SST engages the cingulo-opercular control network to a greater extent, with more pronounced activations in the left anterior insula and bilateral thalamus. Importantly, both tasks also reveal the importance of the anterior insula confirming the role of the SMA [18]. This last was also found to support both action selection and stopping during a voluntary action [19].

In this paper, we investigate the termination process involved during an MI paradigm, and its potential use in BCI applications. We designed a stopping task applied to a MI task by the mean of a clock where subjects are instructed to perform hand MI and stop their MI action as soon as a clock hand is reaching a specific clock tick. The mentioned protocol was tested in offline but also in closed-loop online scenarios.

By doing so, we aim to compare the effect of training a decoder based on the movement-to-rest transition (offset) with the usual decoder trained on the rest-to-movement transition (onset). Here, we hypothesize that (1) the correlates of movement termination during MI tasks are different from the resting state, and (2) that such differences can be captured by a BMI decoder which could hence detect faster and better such transition compare to a usual decoder.

## II. Methods

### 1. Participants

A total of 17 healthy naïve subjects (19-26 years, 8 females, right-hand dominant) participated in the experiment. The study was approved by the Cantonal Committee of Vaud, Switzerland for ethics in human research (CER-VD) and subjects gave their written permission and signed a consent form.

### 2. Offline Protocol

The following protocol was adapted from our previous study [20]. Participants were comfortably seated in front of a PC monitor and asked to perform kinesthetic MI (i.e. imagining the kinesthetic feeling associated with performing a movement) of their right hand while fixating a cross in the middle of a clock shown at the center of the screen. In contrast to our last study, a gauge was added to the design corresponding to a filling green bar and drawn on the clock hand (see Fig. 1). This gauge was gradually increasing with time and subjects were instructed to start their MI once the gauge was visible to them. Once this gauge was filled, the clock hand was activated and initiated its turn. The subject was instructed to maintain his MI task and stop his action (MI termination, MIt) only once the clock hand was overlapping with a red target (offset cue) located on a specific tick. The location of this target was varying in time and was uniformly distributed giving a total duration of MI of 4.35s ± 0.38s (mean ± standard deviation). After terminating their task (MIt), subjects were asked to stay still until the hand clock finishes its turn (4 to 5 s). The total time of a clock hand revolution (MI plus rest) was 8 s. In between trials, a relax period of 5s was introduced between trials. In total, 6 runs of 20 trials were performed by the subject.

**Fig. 1.**
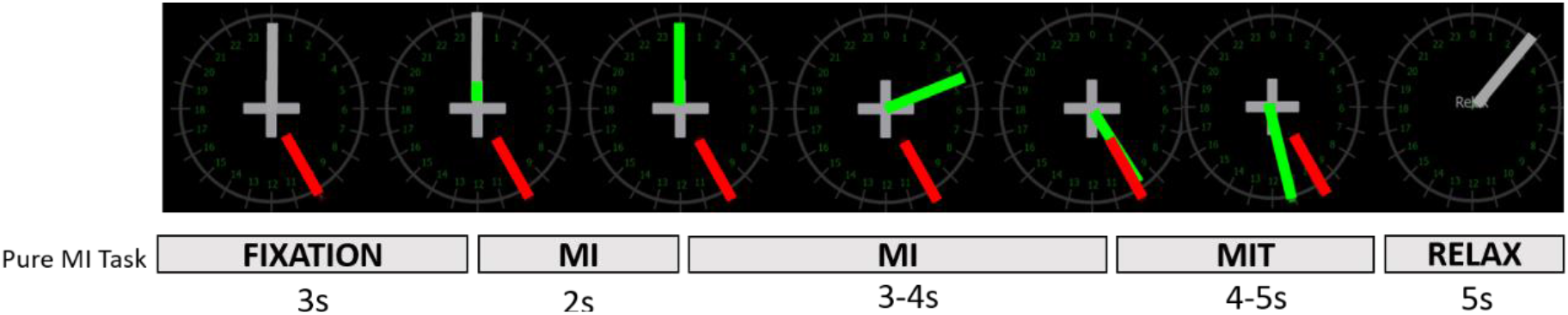
Trial structure during the offline session. During a trial, the subject is asked to continuously look at a fixation cross in the center of the clock. The subject is instructed to stay calm for the first 3 seconds without moving or blinking. Once a gauge represented as a filling green bar is visible, the subject initiates motor imagery (MI) of his right hand. After 2s of MI, the gauge was completed and the clock hand begins his turns around the clock. When the clock hand reaches a target (red bar), the subject should stop motor imagery (MIT) and remain still (no blink or movement) until the clock hand finishes to revolve. A period of 5 seconds following each trial allows the subject to relax.

### 3. Online Protocol

In this part of the protocol, participants were actively controlling the clock through the use of a finite state model with a sequential use of two decoders. Both decoders were trained on the offline data (c.f. Section II-5) to respectively detect the onset and the offset transition. Participants’ task was to initiate the clock hand and to stop later precisely on a target by controlling their MI action. Each subject performed 4 runs of 15 trials each (60 trials in total). Contrary to the offline protocol, the gauge was changing based on the BMI output of the onset decoder as a source of continuous feedback. This continuous feedback was present only for MI onset detection. Here, BMI output corresponded to the integration of the output probabilities of the decoder to each single EEG sample based on an exponential moving average (Eq.1). The smoothing parameter α was set by the operator and kept fixed for the rest of the experiment.

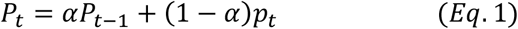

where α is a smoothing parameter, 0 ≤ α ≤ 1, pt is the posterior probability of detecting MI onset and Pt is the probability obtained after smoothing.

Accordingly to the finite state model, once the onset transition was detected, a second decoder was launched and focused on the detection of the MI termination. Here, the detection of MIt was based on an accumulated decision, i.e. counting the number of times the decoder was detecting MIt (pt > 0.5). This parameter was set by the operator and kept fixed for the rest of the experiment as well.

### 4. Recording System

EEG signals were recorded at a sampling frequency of 512 Hz with 16 active surface electrodes placed over the sensorimotor cortex i.e., on positions Fz, FC3, FC1, FCz, FC2, FC4, C3, C1, Cz, C2, C4, CP3, CP1, CPz, CP2, and CP4 according to the international 10/10 system (reference: left earlobe; ground: AFz; g·tec gUSBamp, Guger Technologies OG, Graz, Austria). The amplifier was set with a hardware band-pass filter between 0.01 and 100 Hz (Butterworth 4^th^ order) and a notch filter between 48 Hz and 52 Hz. A common average reference was used on the EEG raw data to enhance the signal-to-noise ratio.

### 5. Analysis of EEG motor correlates

#### a. Time-Frequency Analysis

To study the correlates of motor termination, a spectral analysis was first performed on the following six channels: FCz, C3, C4, Cz, CP3, and CP4. We evaluated the event-related spectral changes (ERSP) with respect to the offset transition [21]. The spectrogram was averaged over subjects and was obtained with the following equation:

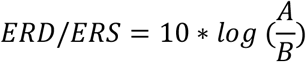

Where A represents the power activity computed with the short-time Fourier transform and for a frequency of interest at a given time and B represents the average power during a baseline interval, chosen between [-2,0] with respect to the onset. The average band power was also calculated by computing the average power over μ and β frequency bands using the same baseline.

#### b. Statistical Analysis

Statistical analysis was performed on the population level to discern significant ERD/ERS patterns with *α* = 0.05 on the ERSP maps. To do so, a non-parametric permutation test (*n =* 5000 permutations) [22] was performed and was corrected for multiple comparisons using max-pixel-based thresholding. A similar test was performed on the averaged band power.

### 6. Offline Classification

#### a. Training

Two decoders (aiming for onset and offset detection) were trained based on the data from the offline experiment protocol. Power spectral densities (PSD) were computed in a 1s-window based on Thomson’s multitaper power spectral density (PSD) estimation from 8 to 30 Hz with a 1 Hz resolution on the 16 channels, yielding a total of 592 features. To assess the classification performance, we performed a nested cross-validation (CV) based on 10 folds where the number of features used to build each model from each fold was fine-tuned. Importantly, these 10-fold were trial-based to avoid any possible overlapping between training and testing sets. For each fold, a diagonal Linear Discriminant Analysis (d-LDA) classifier was trained to distinguish between MI and MI termination processes which were respectively defined between the time intervals [-2, 0] s for MI and [0.5, 2.5] s for MI termination with respect to the offset cue (t = 0). To fine-tune the number of features, inside each fold, we performed an inner 10-fold CV where the number of features was varied between 1 and 50. The optimal number of features was chosen when minimizing the averaged misclassification over the inner CV. Applying a similar process to detect the onset of MI task, a classifier was trained to distinguish between time intervals [-2, 0] s (REST) and [0, 2] s (MI) with respect to the onset (t = 0).

#### b. Classification Metrics

To assess the classification performance, we calculated the accuracy at the sample level over the 10-fold cross-validation. Here, one sample corresponds to PSD features estimated on the 1s-window. The accuracy was defined as the number of correctly classified samples over the total number of samples and was computed for each fold. We estimated the chance threshold at the 95% confidence interval based on the inverse binomial cumulative distribution.

#### c. Pseudo-online classification

A pseudo-online (PO) analysis was performed on the data to further study the behavior of our offset decoder on the offset transition. In this analysis, the classifier trained to detect the offset transition was tested in the time interval [-3, 4] s with respect to the offset cue. During this time interval, the likelihood for each class was calculated from the decoder on samples computed with a 1s-window shifted every 62.5 ms. Using this decoder, we measured the decoding latency, which we defined as the time when the average posterior probabilities over trials were crossing the chance threshold.

#### d. Comparison of decoding approach

For the pure MI task we compare the respective behavior of each decoder when applied in a pseudo-online analysis to detect its own transition (onset PO, offset PO). Through this comparison, we compared each decoder type regarding their accuracies to detect MI termination when tested on the interval s [-2, 0] s (MI) and [0.5, 2.5] s (MI termination) with respect to the offset (t = 0) and compare their latency via a pseudo-online analysis in the time interval [-3, 4] s with respect to the offset cue. Importantly, using the onset decoder, the decoding likelihood to detect MI was inverted to obtain the decoding likelihood to detect the rest state while for the offset decoder, the decoding likelihood to detect MI termination was kept.

#### e. Random Forrest Classification

A Random Forest for the three-class problem was also tested on a 10-Fold cross-validation using the same features as before (see Section 5.a) for the three following classes: REST, MI, MI termination. The classifier was trained with 1000 trees with a maximal depth of 5. The features were selected by Random Forest that automatically ranked features based on how they improve the purity of the node. Similarly to Section 5.c, we performed a pseudo-online analysis where we reported the decoding likelihood for each class when an RF-based classifier was tested on a time window [-3, 4]s with respect to either the onset or the offset transition. Such alternative to decode both onset and offset transition could bring benefits to current decoding process as it would decrease the false positive rate. This alternative approach was compared with our previous decoding approach (diagonal LDA + Fisher Score, d-LDA + FS) for the same 3-class problem.

#### f. Statistical Analysis

To compare the classification accuracies between the onset and offset classifiers, a paired t-test was used on the mean accuracies per subject. Similarly, a paired t-test was also performed on the average detection over trials. Finally, the average fisher score was also calculated over μ and β bands and averaged over subjects, a paired t-test was also performed on these data.

### 7. Post-hoc analysis on online data

Analyzing the online result, three different type of detection were defined based on the time of detection (early detection: < -1.5s, correct detection: [-1.5, 1.5] s, late detection: > 1.5s). To characterize each type, we computed for each type of trials, the fisher score between the sample [-1, 0] s and [-3, -2] s samples.

## III. Results

### 1. Neurophysiological signature of MI termination is different from the resting state

Fig 2 shows the grand average across subjects recorded during pure MI task as well as the power averaged over the µ band [10-14] Hz and β band [20-30]. For each band, we reported the results respectively in blue for the µ band and in red for the β band. When plotting the band powers, we additionally drew horizontal lines to show when an ERS has been found significant based on non-parametric permutation test (α = 0.05, repeated measure t-tests based on Wilcoxon rank-sum test, FDR corrected for multiple comparisons). The channels were grouped according to their location on the scalp (central, contralateral, and ipsilateral channels). A significant ERS can be observed in contralateral channels (C3, CP3) as well as for central channel Cz. This increase is mainly characterized in the β band and start 1s after MI-STOP indicating the presence of β rebound. An increase of power in central channels FCz and Cz can also be observed.

**Fig 2.**
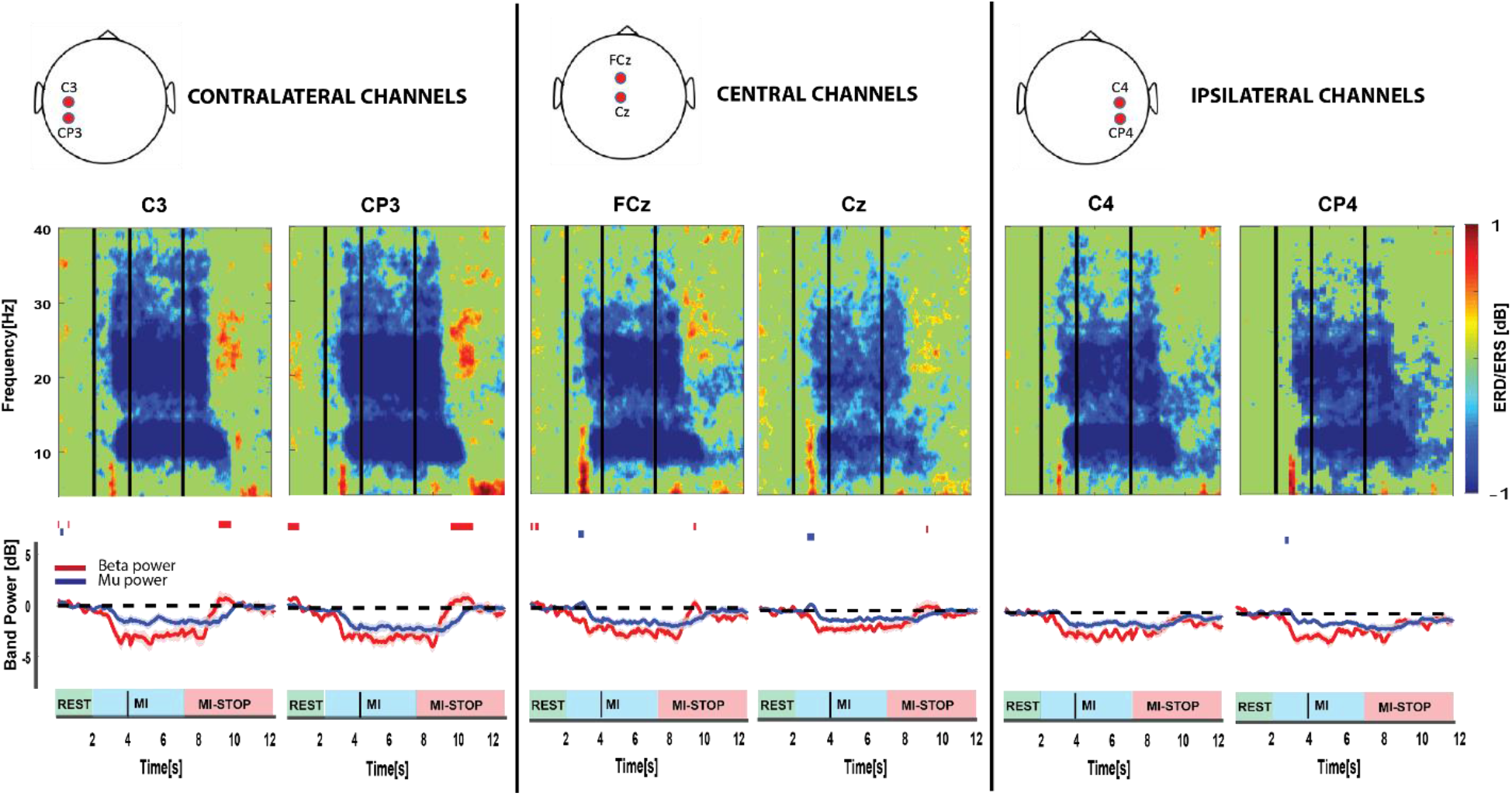
Grand average spectrogram across subjects and averaged over all trials during the offline session. The vertical lines correspond respectively to the different event in the protocol: starting cue for MI (1^st^ line), the activation of the clock (2^nd^ line), and finally, the stopping cue for MI (3^rd^ line). ERD/ERS were computed with a logarithmic scale using a baseline period [-2 0] s with respect to the starting cue. Below the spectrogram, band power of μ band (in blue) and β band (in red) for each channel averaged over all the trials are shown. The time origin corresponds to the offset. Red horizontal lines on top indicate periods of significant difference for the β band based on the baseline period for its corresponding band (α = 0.05, repeated measure t-tests based on Wilcoxon rank-sum test, FDR corrected for multiple comparisons).

### 2. BMI for MI termination

We first built two classifiers to detect each motor state transition, namely the rest-to-movement (onset) transition and the movement-to-rest (offset) transition. After that, we evaluated the performances of both onset and offset decoders. For each subject, we reported in Table 1 the mean accuracies as well as their standard deviation of the 2-class d-LDA averaged over nested cross-validation and across subjects. On one hand, when decoding the onset, an offline average accuracy was achieved at the sample-based level of 68.0% ± 6.4% (mean ± std). On the other hand, in the case of the offset, an average accuracy of 69.7% ± 7.0% was obtained. Importantly, both classifiers decode above the statistical chance threshold (54.17%) for every subject. We reported in Table 1 for each subject the latency of their decoder for each transition. In general, one can notice that the onset latency is smaller than the offset latency (paired t-test, *t(16)* = 4.46, *p* = 3.9e-04 < 0.001 ***) while the accuracies were not found significantly different (paired t-test, *t(16)* = 1.05, *p* = 0.31 > 0.05).

**Table 1.**
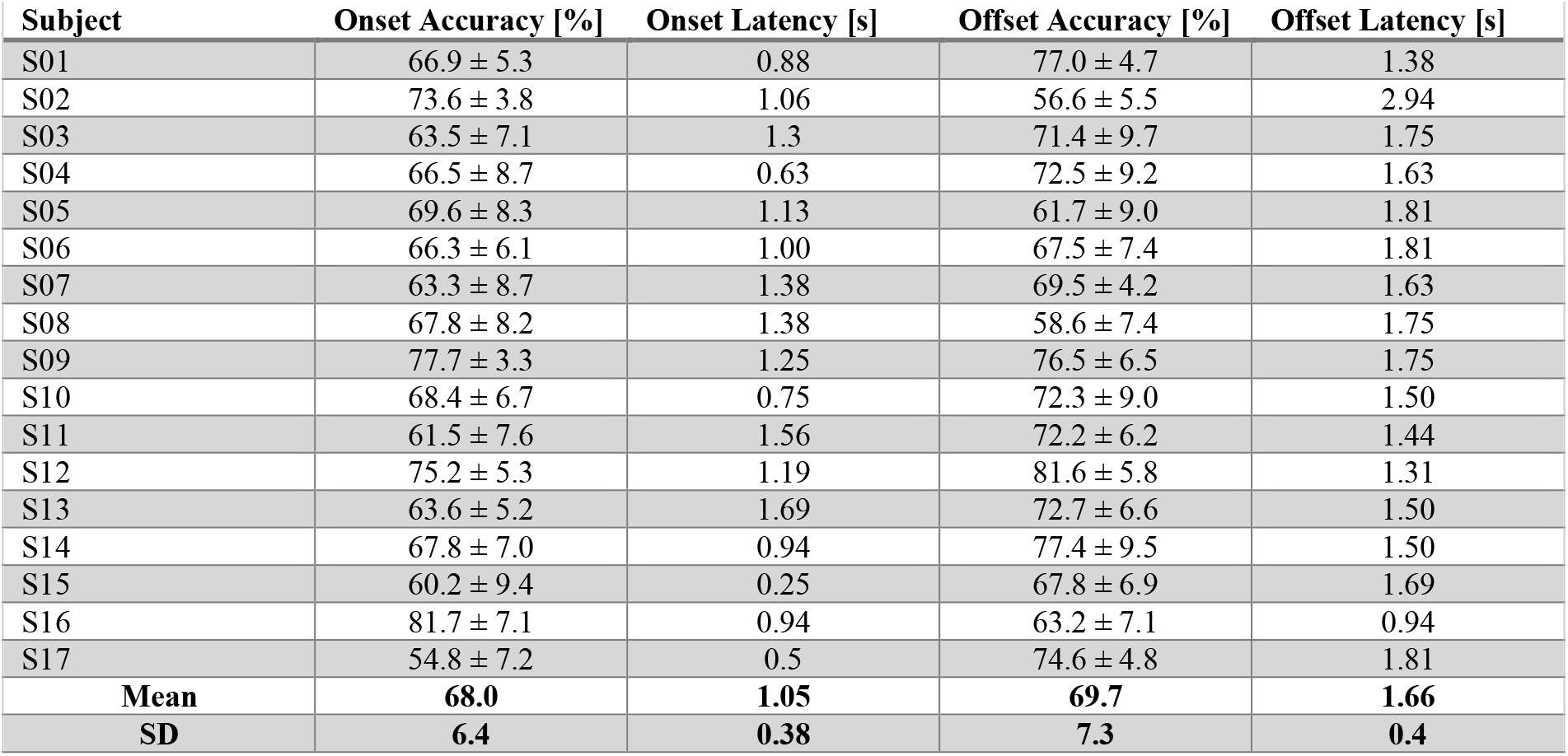
Classification accuracy for motor state transition decoding. The accuracies and detection latencies was reported for each subject as well in grand average with their mean and standard deviation (SD) for both onset and offset decoding.

| Subject | Onset Accuracy [%] | Onset Latency [s] | Offset Accuracy [%] | Offset Latency [s] |
| --- | --- | --- | --- | --- |
| S01 | 66.9 ± 5.3 | 0.88 | 77.0 ± 4.7 | 1.38 |
| S02 | 73.6 ± 3.8 | 1.06 | 56.6 ± 5.5 | 2.94 |
| S03 | 63.5 ± 7.1 | 1.3 | 71.4 ± 9.7 | 1.75 |
| S04 | 66.5 ± 8.7 | 0.63 | 72.5 ± 9.2 | 1.63 |
| S05 | 69.6 ± 8.3 | 1.13 | 61.7 ± 9.0 | 1.81 |
| S06 | 66.3 ± 6.1 | 1.00 | 67.5 ± 7.4 | 1.81 |
| S07 | 63.3 ± 8.7 | 1.38 | 69.5 ± 4.2 | 1.63 |
| S08 | 67.8 ± 8.2 | 1.38 | 58.6 ± 7.4 | 1.75 |
| S09 | 77.7 ± 3.3 | 1.25 | 76.5 ± 6.5 | 1.75 |
| S10 | 68.4 ± 6.7 | 0.75 | 72.3 ± 9.0 | 1.50 |
| S11 | 61.5 ± 7.6 | 1.56 | 72.2 ± 6.2 | 1.44 |
| S12 | 75.2 ± 5.3 | 1.19 | 81.6 ± 5.8 | 1.31 |
| S13 | 63.6 ± 5.2 | 1.69 | 72.7 ± 6.6 | 1.50 |
| S14 | 67.8 ± 7.0 | 0.94 | 77.4 ± 9.5 | 1.50 |
| S15 | 60.2 ± 9.4 | 0.25 | 67.8 ± 6.9 | 1.69 |
| S16 | 81.7 ± 7.1 | 0.94 | 63.2 ± 7.1 | 0.94 |
| S17 | 54.8 ± 7.2 | 0.5 | 74.6 ± 4.8 | 1.81 |
| **Mean** | **68.0** | **1.05** | **69.7** | **1.66** |
| **SD** | **6.4** | **0.38** | **7.3** | **0.4** |

To study the behavior of each classifier during both transitions, we performed a pseudo-online analysis where each decoder was applied in the time intervals [-4, 4] s with respect to the corresponding transition. We can observe a decoding likelihood of MI initiation that seems to reach a plateau which can be interpreted as the detection of sustained MI (Fig. 2A) while for MI termination, the decoding likelihood reaches a peak around 2s decreasing rapidly after indicating the detection of a fast and transient state (Fig. 2B).

**Fig 2.**
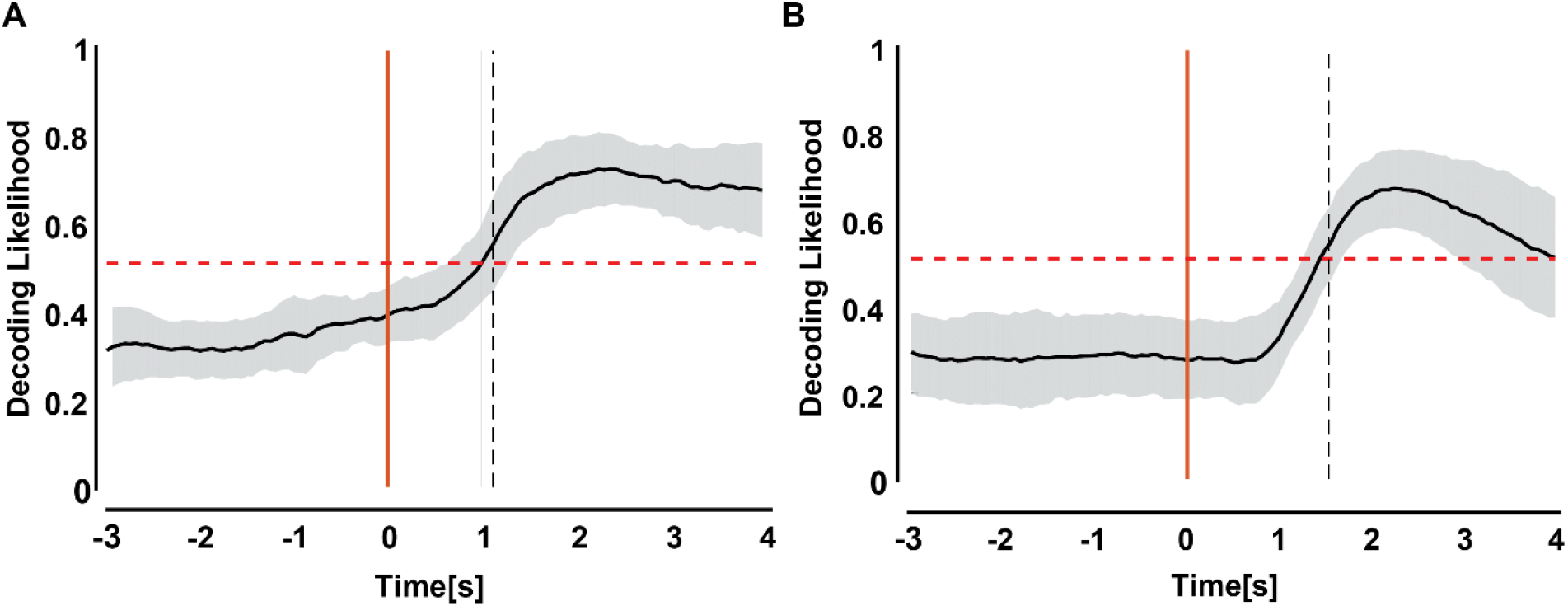
Grand Average pseudo-online analysis of each decoder on their transition. The likelihood of both decoders averaged over subject is shown over time. A. Pseudo-online decoding of onset decoder applied on the initiation transition (t=0) of MI. b) Pseudo-online decoding of offset decoder applied on the termination transition (t=0) of MI. The standard deviation is shown at every time point in grey. The chance level is represented with a horizontal dashed red line.

After building both decoders, we compared each decoder regarding their performances to detect the offset. In Figure 3A, we show the Fisher Score for each feature averaged over subjects and mapped it in 2D (channels x frequencies). On the grand average, when looking at the selected features during the decoding process, both transitions show relevant information on similar spatial locations, i.e. contralateral channels (C3, CP3). However, the features selected for each decoder differ in frequency bands with an onset decoder primarily based on features in the µ band and an offset decoder relying principally on features in the β band. Comparing the average fisher score for both decoder, we found that μ band was statistically higher for the onset decoder overall channels (paired t-test, *t(16) = 3*.*45, p* = 0.0033 < 0.01 **). No significant difference was found for β band (paired t-test, *t(16)* = 0.46, *p* = 0.65 > 0.05).

**Fig 3.**
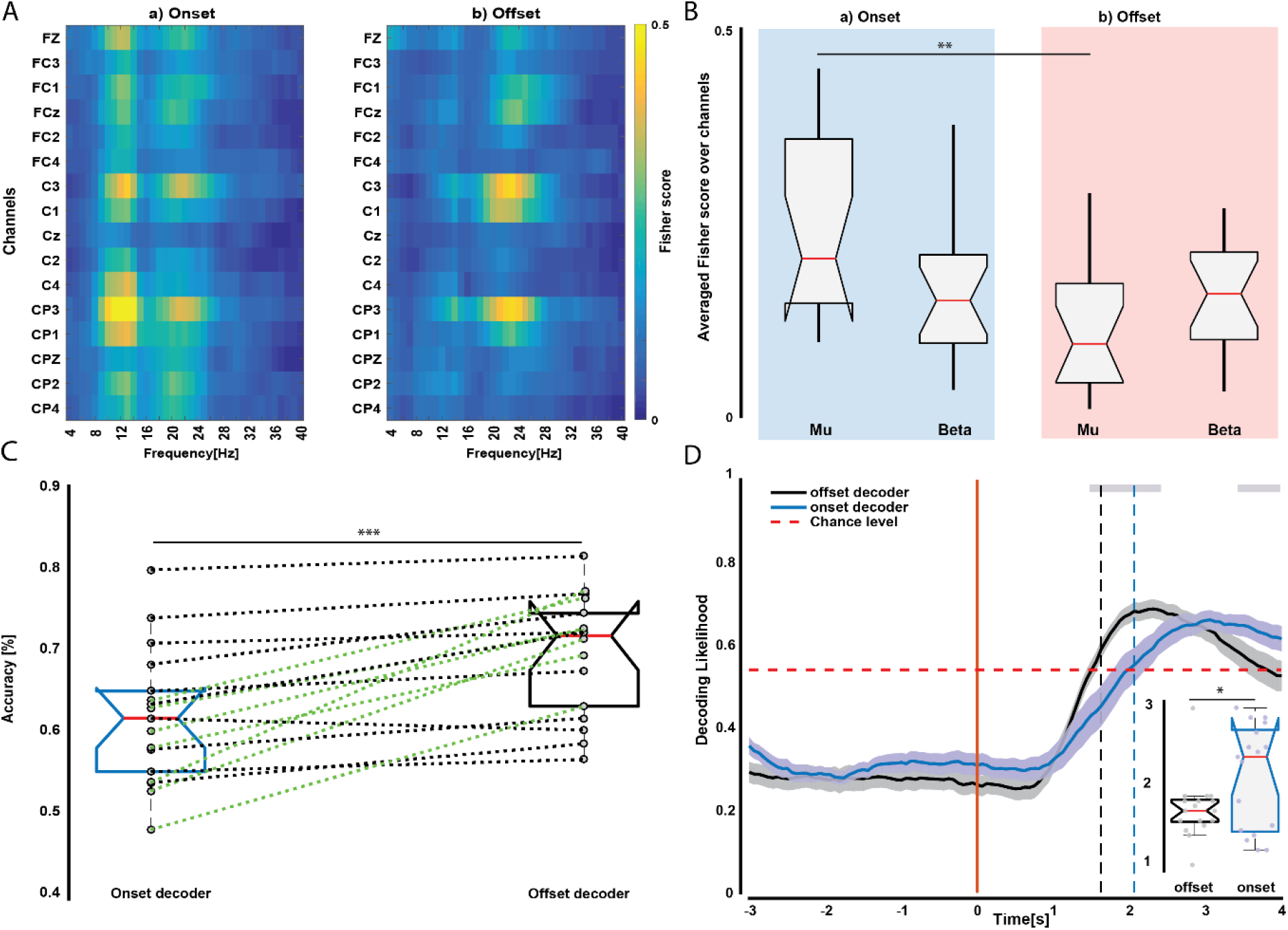
Comparison between both decoding. A. Map of selected features averaged over all subjects for each decoding approach. B. Average Fisher score over all channels for μ ([8-12] Hz) and β ([13-30] Hz) bands. C. Classification accuracies of motor termination by both decoders are reported using boxplot with each dash line representing a subject (green is showing a significant difference of accuracies). D. Pseudo-online analysis of the decoding termination averaged over subjects for each decoder. The solid vertical line corresponds to the offset transition and the horizontal dash lines correspond to the time when the decoding likelihood is reaching the chance threshold. Grey rectangles on the top of the plot show a significant period where the onset and offset decoder show a significant difference. These periods were found based on statistical analysis (non-parametric Wilcoxon rank-sum test, α = 0.05). Boxplot for latency detection was also reported on the bottom right corner for both decoding.

Then, we compared the decoders regarding their performances to detect the offset transition. In Fig 3C, we reported the accuracies for each subject on the nested 10-fold CV showing an increase of accuracy when applying an offset-specific decoder on the offset transition (69.7 % ± 7.3% for the offset decoder against 61.7% ± 7.3% for the onset decoder). This increase was statistically tested (paired t-test, *t(16)* = 4.75, *p* = 2.1e-4 < 0.001 ***). All subjects showed an increase of accuracy using the offset decoder which was significant for 7 of them. Finally comparing both decoders in a pseudo-online analysis, we show that the offset decoder can detect the transition faster with an earlier detection occurs at 1.58s ± 0.36s while using the onset decoder, the detection occurs at 1.97s ± 0.67s (paired t-test, *t(16)* = 2.67, *p* = 0.017 < 0.05 *).

Comparing the decoding likelihood of both decoders over the pseudo-online analysis, we also reported a significant difference in the dynamic of the probabilities (non-parametric Wilcoxon rank-sum test, α = 0.05). This difference can be explained as our offset decoder detects the β rebound since our decoder was specifically trained on the time interval which exhibits the β rebound. Then our decoder can detect this pattern of motor termination while the onset decoder –not specifically trained to recognize such pattern-was detecting only the rest period happening after the β rebound hence explaining the delay in the offset detection.

### 3. Online Protocol Results

During the online session, subjects were instructed to initiate the clock hand by performing MI, and then, to stop the same clock hand by stopping their MI. In Fig 5, we reported the results of this experiment showing that on the grand average subjects learn to stop the clock hand effectively. Putting all the trials together, we found that on the grand average we found that the BMI was stopping the clock hand with a median of 0.62s [Q1: 0.89s, Q3: 1.87s] (Fig. 5A). Following these results, we reported the fisher score before the detection made by the BMI. In results, visual inspection shows that the decision taken from BCI on the correct detection interval relies more on β rebound on contralateral channels (C3, CP3) as well as central channel FCz while the late detection will rely additionally on μ features with more contra lateralization (Fig. 5B). These results support our previous results and reinforce our previous findings on the difference between BCI correlated of REST and MI termination. Moreover, they show that β rebound allows sharper detection allowing our decoding to be more accurate in time.

### 4. Three Class Model

In this section, we investigated the feasibility to decode through a unique decoder both onset and offset transition. We reported the classification accuracies of both approaches in Table 2 for each subject. Results indicate that RF can decode simultaneously better both motor state transitions. Indeed, an average accuracy of 48.9% ± 4.4% was found for d-LDA+FS approach while an average accuracy of 58.2% ± 5.9% was found for RF corresponding to an almost 10% significant increase (paired t-test, *t(16)* = 10.570, *p* = 1.3e-8 < 0.001 ***). Confusion matrix averaged over subjects was also reported for RF in Fig. 4B showing that RF can reliably decode each state (REST, MI, MIt) above chance level (> 0.3). Similar to the previous analysis, we first looked at the features importance of our classifier. In this case, RF automatically performs feature selection. For each subject, we computed the features importance mapping and averaged it over subjects. In Fig. 4a we show the results of these mapping averaged over subjects. Similar to Fig. 3A contralateral channels (C3, CP3) are primarily selected by the Random Forest with both μ and β bands carrying relevant information to discriminate the three different classes. Finally, we also performed a pseudo-online analysis on both onset and offset transition using our RF classifier. Results show a clear period devoted to each of these transitions. Importantly, during the offset transition, the distinction between MIt and Rest is captured by our decoder. By looking at the dynamic of the decoding likelihood, we can strongly conclude that RF relies on β rebound for detecting MIt. Once this rebound is not detected, the rest state follows.

**Table 2.** Classification accuracy for the three-class problem. Mean accuracies and their standard deviation for each classifier was reported in both methods (diagonal LDA + Fisher Score, Random Forest Classification). This was done for each subject as well in grand average with their mean and standard deviation (SD).

| Subject | d-LDA + FS | RF |
| --- | --- | --- |
| S01 | $53.9 \pm 6.4$ | $62.4 \pm 7.8$ |
| S02 | $45.3 \pm 5.1$ | $48.9 \pm 6.0$ |
| S03 | $54.4 \pm 6.8$ | $64.5 \pm 10.4$ |
| S04 | $51.8 \pm 8.0$ | $57.1 \pm 6.2$ |
| S05 | $43.1 \pm 6.5$ | $48.2 \pm 5.0$ |
| S06 | $46.0 \pm 5.3$ | $60.4 \pm 5.7$ |
| S07 | $44.8 \pm 7.3$ | $53.9 \pm 4.5$ |
| S08 | $43.3 \pm 6.1$ | $51.4 \pm 6.0$ |
| S09 | $51.0 \pm 4.5$ | $67.3 \pm 3.2$ |
| S10 | $50.2 \pm 5.6$ | $56.7 \pm 6.8$ |
| S11 | $50.8 \pm 5.1$ | $59.8 \pm 5.4$ |
| S12 | $57.3 \pm 6.9$ | $65.8 \pm 5.8$ |
| S13 | $44.2 \pm 5.4$ | $57.0 \pm 4.6$ |
| S14 | $53.4 \pm 9.0$ | $66.4 \pm 5.6$ |
| S15 | $45.3 \pm 8.2$ | $59.5 \pm 5.7$ |
| S16 | $48.3 \pm 2.9$ | $55.2 \pm 6.7$ |
| S17 | $48.3 \pm 5.3$ | $55.2 \pm 5.9$ |
| **GA** | **$48.9 \pm 4.4$** | **$58.2 \pm 5.9$** |

**Fig 4.**
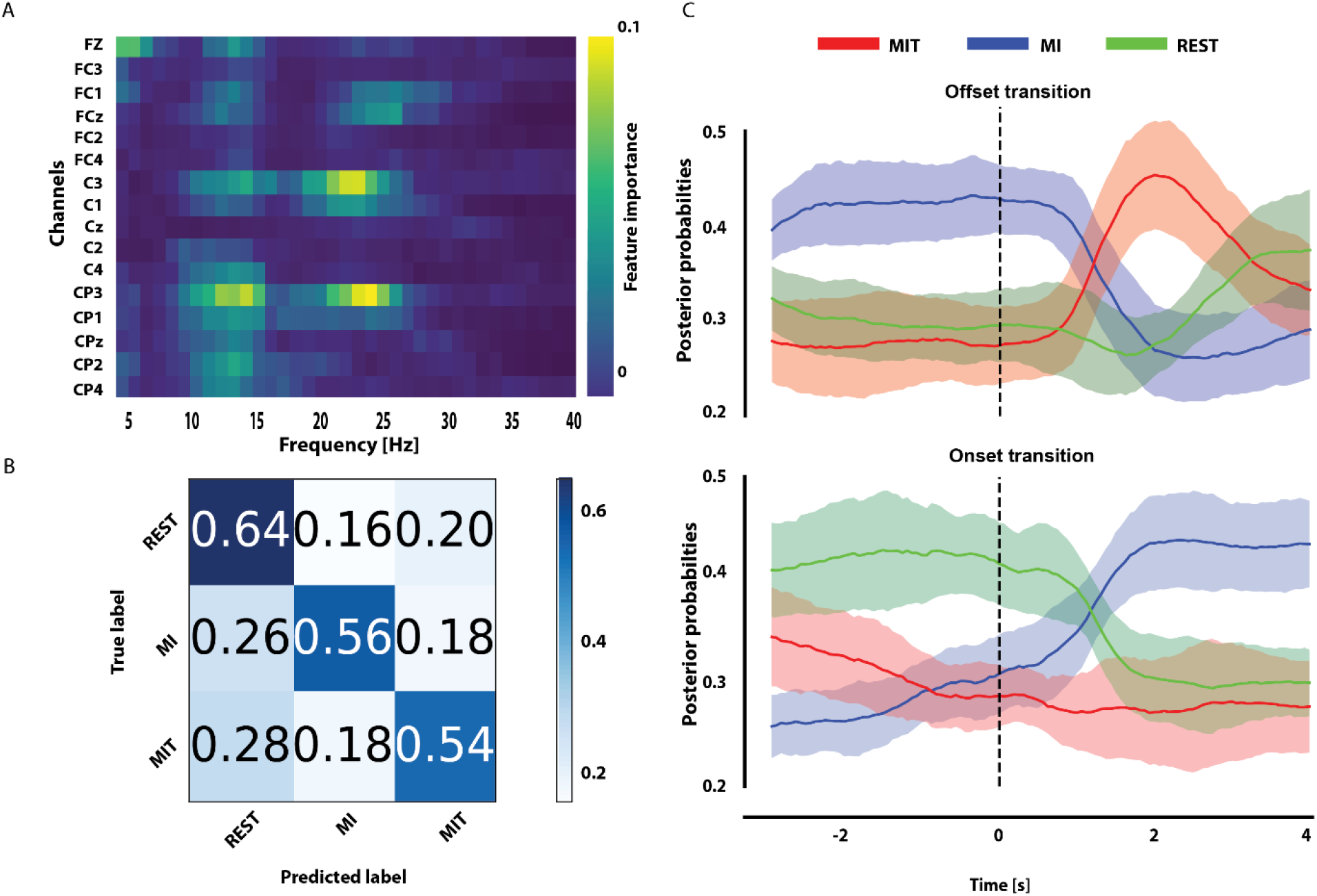
Random Forest for decoding motor state transition. Three class problems with Random Forest for decoding motor state transition. A. Map of features importance ranked by Random Forest classifier and averaged over all subjects. B. Confusion matrix averaged over all subjects. C. Pseudo-online analysis using Random Forest on both offset (top) and onset transition (bottom). Both transitions are represented at t=0 with a black dash line.

**Fig. 5.**
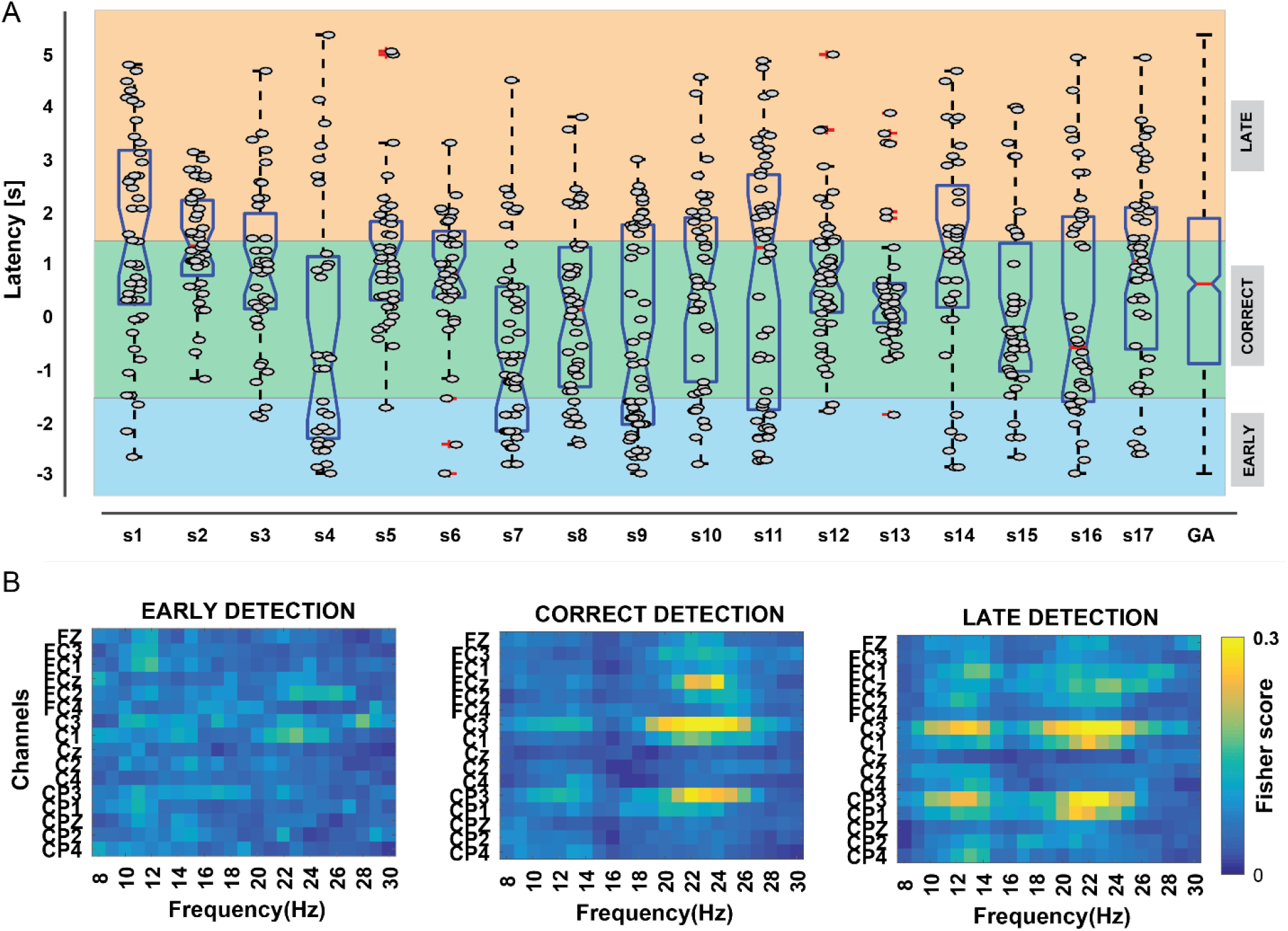
Performances during the closed-loop experiment. A) Distribution of latency trials for each subject. Each point corresponds to the latency of one trial and was calculated by computing the difference between the time when the participant is supposed to stop and when the clock hand was stopped by BCI. Boxplots illustrate the distribution and the median latency for every subject as well as on the grand average. Colored areas represent the intervals of different detection types (early detection: < -1.5s [blue], correct detection: [-1.5, 1.5] s, late detection: > 1.5s [orange]). B) Fisher score map averaged over all subjects. For each detection type (early, correct, late), we computed for the corresponding trials the fisher score between [-1 0] s and [-3, -2] s samples. Fisher scores are shown for the features (channels x frequencies). Higher values (i.e., yellow color) indicate highly informative features while blue colors indicate less informative features. The scores were normalized for each subject using min-max scaling.

## IV. Discussion

In this paper, our goal was to investigate the differences between the movement termination and the resting state process involved during a motor imagery task from the neurophysiological and brain-machine interface point of view. To do this, we proposed that (1) the correlates of movement termination during a MI task are different from the resting state, and (2) that such differences can be captured by a BMI decoder.

### 1. Neurophysiological differences between MI termination and Rest

The EEG signals were first analyzed via a time-frequency analysis which was applied to the three main periods of interest: the resting state period, the MI period, and the MI termination. As expected, the analysis revealed first the MI correlates, i.e. a decrease of power in μ and β band (μ-ERD and β-ERD). On the opposite, the MI termination is predominantly characterized by an increase of power in the β band (β-ERS, β rebound). Such synchronization, consistent with the literature, appears around 1s after MI end and is located mainly over the sensorimotor cortex on contralateral channels. Although less important, an increase of power in FCz can also be observed, which is consistent with the proposed the roles of the premotor supplementary area (preSMA) in the process of stopping a voluntary action [16], [17].

Although this characterization of the termination process involved during a MI task is consistent with the literature, our previous experiments also showed additional correlates. Indeed, in our previous experiment [20], the presence of a μ ERS was found to be predominant and more reliable over subjects while here a β ERS is predominant. This strong component of rest was appearing shortly after the termination of MI and was observed until the end of the trial making such correlates strong features for detecting the end of Motor Imagery. Such differences can be explained mainly by the instruction given to the subjects. First, in the previous experiment, subjects were using both hands while here, the use of the dominant hand was asked. Second, in this previous experiment, subjects were asked to perform a repetitive kinesthetic MI while here, subjects were instructed to do continuous MI. These differences could then explain such conflicting results. While the first difference is dominantly influencing the lateralization of MI termination correlates [23], [24], the second can induce important changes in these correlates [25]. Indeed, while comparing brief and continuous MI, Nam and his colleagues found a duration effect on MI termination correlates. Importantly they showed that μ power was returning to the baseline more quickly in continuous MI task while on the opposite Beta powers were returning more quickly during the brief MI [26]. This finding supports our results and explains the absence of µ ERS in this study. Importantly, this implies as well that the termination process might differ based on the type of MI which makes it then a critical factor to take into account for detecting MI termination and the creation of BMI models.

### 2. Evidence for the use of a different model to detect MI termination

Most MI-based BMI studies aim to detect MI as a sustained activity. Here, we investigated two different approaches: (1) a finite state model composed of two classifiers to detect both motor state transitions i.e. rest-to-movement (onset) and movement-to-rest (offset) transition, and (2) a three-class Random Forest classifier trained to discriminate between the three classes together (MI, REST, and MIt).

Using a finite state model, we trained two decoders on the onset and offset transition. By training a specific decoder trained to detect MI termination, we were able to outperform the standardized BMI decoder based on MI and Rest classes. Our decoder shows faster detection and higher accuracy. Such improvement can be mainly explained by the use of β features that are associated with the post-movement rebound phenomenon. Since such rebound appears earlier than the baseline return of μ power, these features are more suitable to detect MI termination. On the opposite, a standardized BMI for onset detection will principally rely on μ features and will wait for MI correlates to collapse delaying the detection of MI termination. Additionally, using the finite state model in a closed-loop experiment, a predominance of β features can also be noticed for the trials detected within the correct time intervals while for the late detection additional features in μ band and associated with an idling process [13], [27], [28] can be observed reinforcing the importance of β features for decoding MI termination while being timely accurate. Nonetheless the performance of a finite-state based decoding will be bounded by the performance of the MI onset decoder. Using a 3-class model, a difference between resting state and MI termination was also made by the Random Forest classifier. This classifier was trained to discriminate between the three classes (MI, REST, and MIt) altogether by notably outperforming the LDA classifier by almost 10%. Such differences can easily be explained by the approach used by different classifiers where the multi-class LDA we used was partitioning the different classes to have one class against the rest resulting in 3 classifiers and combining the results while Random Forest was considering the three classifiers together. Hence, Random Forest shows evidence that it is possible to discriminate between the three different classes. More importantly, such a decoder can differentiate between rest and MI termination after MI action showing a transient state for MI termination quickly followed by a durable resting state. More RFs, being a fusion of weak classifiers, are also more robust when dealing with less samples.

Altogether, these findings contradict the current belief to use asynchronous onset decoding on [9] for the detection of the termination process during MI. The two models can be used as alternatives solutions of the usual approach aiming to detect the onset transition. More, these alternatives provide more accurate and more precise detection of the termination process involved during a motor imagery task.

